# Peptide and protein alphavirus antigens for broad spectrum vaccine design

**DOI:** 10.1101/2022.05.26.493643

**Authors:** Catherine H. Schein, Grace Rafael, Wendy S. Baker, Elizabeth S. Anaya, Jurgen G. Schmidt, Scott C. Weaver, Surendra Negi, Werner Braun

## Abstract

Vaccines based on proteins and peptides may be safer and more broad-spectrum than other approaches Physicochemical property consensus (PCP_con_) alphavirus antigens from the B-domain of the E2 envelope protein were designed and synthesized recombinantly. Those based on individual species (eastern or Venezuelan equine encephalitis (EEEVcon, VEEVcon), or chikungunya (CHIKVcon) viruses generated species-specific antibodies. Peptides designed to surface exposed areas of the E2-A-domain were added to the inocula to provide neutralizing antibodies against CHIKV. EVC_con_, based on the three different alphavirus species, combined with E2-A-domain peptides from AllAV, a PCPcon of 24 diverse alphavirus, generated broad spectrum antibodies. The abs in the sera bound and neutralized diverse alphaviruses with less than 35% amino acid identity to each other. These included VEEV and its relative Mucambo virus, EEEV and the related Madariaga virus, and CHIKV strain 181/25. Further understanding of the role of coordinated mutations in the envelope proteins may yield a single, protein and peptide vaccine against all alphaviruses.

## Introduction

The alphaviruses (AV) are a diverse genus of plus-strand RNA species that are organized into antigenic complexes. While primarily infecting wild and domesticated animals, some can cause encephalitic (eastern and Venezuelan equine encephalitis viruses (EEEV and VEEV, respectively)) and others arthralgic (chikungunya, CHIKV) infections in humans. Both EEEV [1] and VEEV [2-4] cause livestock outbreaks with 80% or higher case-fatality, which have resulted in the death of thousands of horses, pigs [5] and birds. Although human cases of EEEV are rare, they have very high mortality. Giving further cause for concern, EEEV is spread by birds in a fashion similar to West Nile virus, suggesting that it could expand its range [6].

Despite their potential for rapid spread, as has been illustrated by worldwide CHIKV outbreaks [7, 8], there are currently no approved vaccines or specific treatments for human use against any alphavirus. However, there are many experimental vaccines against individual alphaviruses in various stages of preclinical and clinical testing [9-24]. There are also ongoing efforts to generate a trivalent vaccine by combining antigens of EEEV, VEEV and WEEV [25]. However, producing combination vaccines increases production costs while immune interference can prevent maximum response to some or all of the components [26, 27]. Thus, our goal is to design a single vaccine that would provide protection against all AV, and overcome species specificity. Vaccines based on synthetic peptides and proteins require no mammalian cell culture and can be formulated/adjuvanted for different delivery systems[28, 29].

As with other RNA viruses [30-32], variation in the sequences of AV complicates the development of treatments with antiviral drugs, vaccines or monoclonal antibody preparations. Basing mAb treatments on single sequences can also lead to rapid selection of resistant mutants [33, 34]. The specificity of the immune response implied by studies with mAbs has been interpreted to mean that protection against different alphaviruses can only be induced by mixing antigens, virus-like particles or whole viruses from different species [35, 36] or with rearranged genomes. While US isolates of EEEV from many different hosts show little variation [1](Fig. S1), VEEV is more heterogeneous with up to 20% (and higher for mosquito isolates) amino acid sequence diversity [2, 37]. Both EEEV and VEEV differ considerably from each other, and both vary even more from CHIKV and other arthralgic AV strains, by over 60% (Fig. 1). Yet, we have observed that an attenuated VEEV/IRES vaccine can induce partial protection against Madariaga virus (MADV), a member of the EEEV antigenic complex [38].

**Figure 1:**
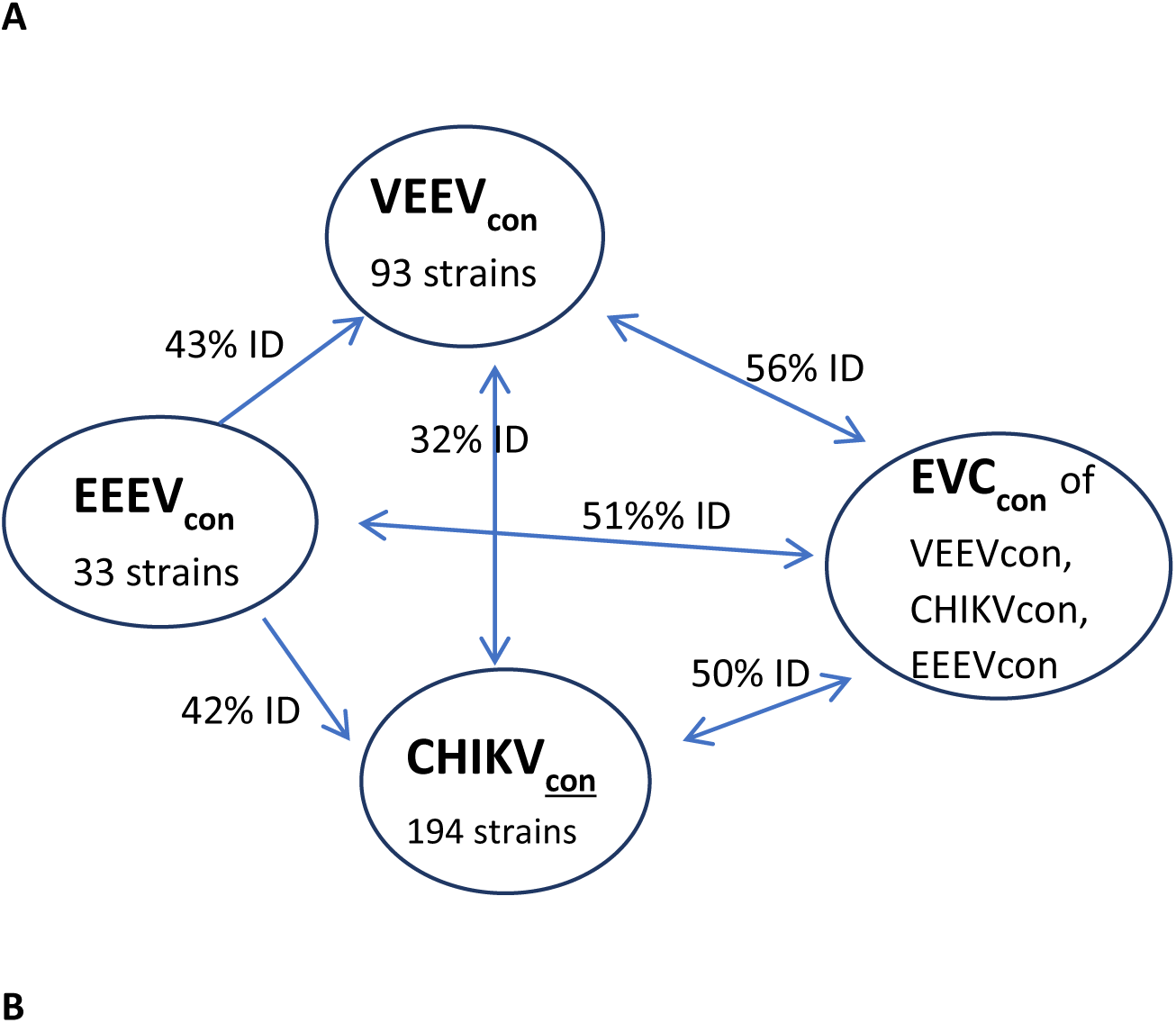
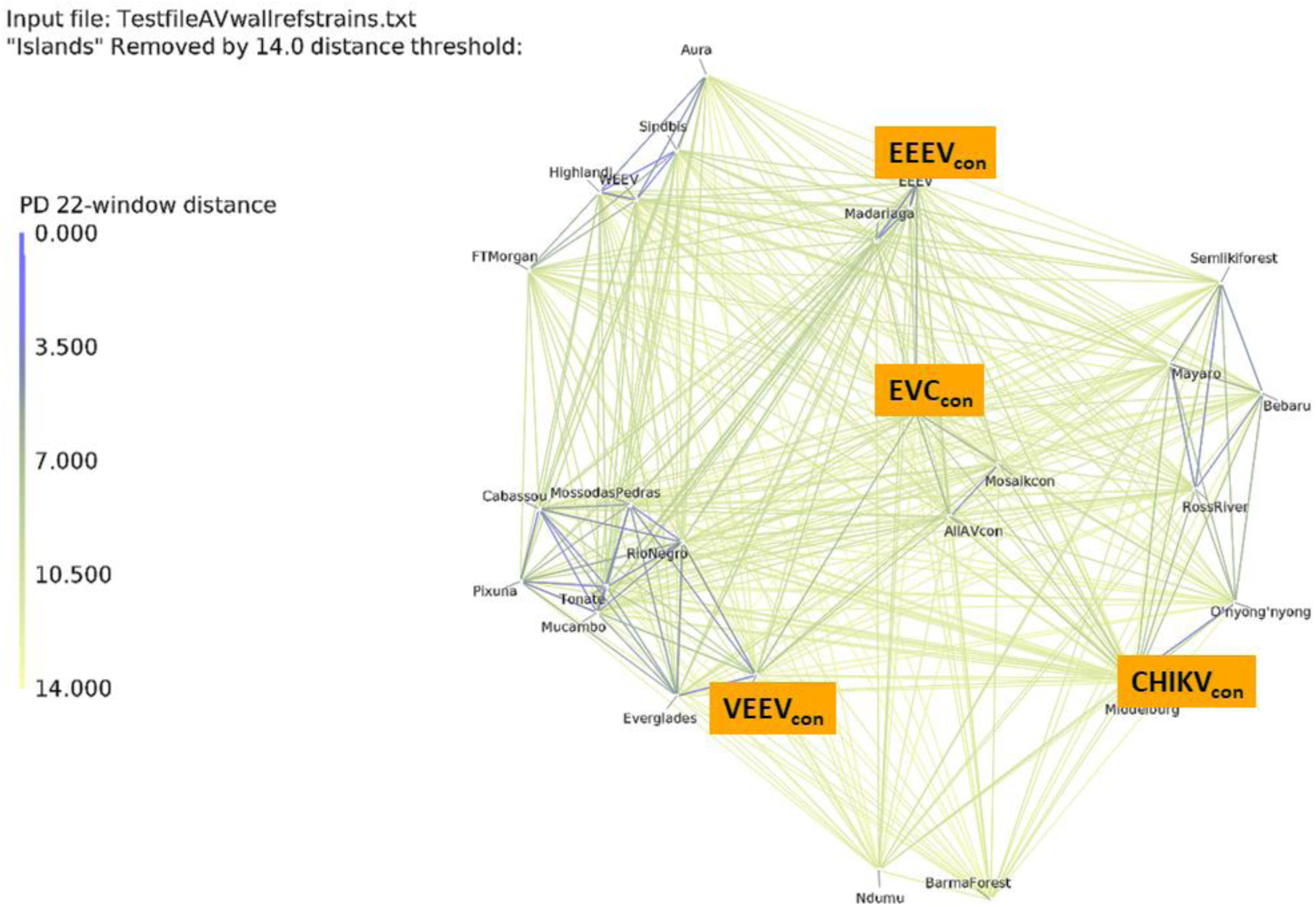
Design of species-specific and broad spectrum (EVC_con_) PCP_con_ antigens. A) The three proteins on the left were calculated using the PCPcon program from individual unique sequences of VEEV, EEEV or CHIKV collected from the VIPR database. Then the three sequences were used to calculate EVC_con_. B) PD-graph[56] of alphavirus reference and PCP_con_ sequences (highlighted in orange, overlaying their graphed position) used in this work show that EVC_con_ and previously described sequences AllAV and Mosaik_con_[45] are true PCP-mean representatives of this diverse family. Lower property distance (PD) between viruses, as specified by the line color and thickness in the Y-axis of the PD-graph, correlates with higher similarity [56, 74].

Other recent studies have indicated that some mAbs can protect against both WEEV and EEEV in murine aerosol challenge [9]. Further, mAbs to the highly conserved fusion loop of E1 (which lies below the B-region of the E2 protein in the intact virus) have cross-neutralizing activity [39], confirming that diverse AV have a similar mechanisms of cell entry. Humans infected with EEEV can produce antibodies that also limit infection with VEEV or CHIKV [39], while those infected with CHIKV can produce antibodies that protect against multiple alphaviruses [40]. Most of the neutralizing mAbs against CHIKV and EEEV that have been structurally characterized bind to surface-exposed regions of the A and B domain of the E2 protein [41-44]. We are thus designing immunogens based on variable, surface-exposed regions of the E2 protein that will induce a promiscuous response and offer more robust protection against variants. We previously showed that PCPcon spike proteins based on the individual antigenic complexes, VEEV_con_, or CHIKV_con_, are recognized exclusively by antisera generated during infection with their cognate viruses, as would be expected based on their low sequence identity (<35%). In contrast, antisera generated during either VEEV or CHIKV infection recognize the PCP_con_ proteins, AllAV and Mosaik_con_, which were calculated based on the sequences of VEEV_con_, CHIKV_con_ and many additional AV reference strains [45], including other pathogens such as Ross River virus [46].

Here, we extended our protein repertoire to include EEEV_con_, and designed EVC_con_, a completely computationally derived protein based on EEEV_con,_ VEEV_con_, and CHIKV_con_. Both proteins have intermediate identity to both VEEV_con_ and CHIKV_con_ (Fig. 1a). We show that while recombinant EEEV_con_, VEEV_con_ and CHIKV_con_ generate specific neutralizing antibodies, their consensus, EVC_con_, generates a promiscuous immune response, able to neutralize representatives of all three species. Thus, this single protein antigen can generate antibodies against very diverse alphaviruses.

## Materials and Methods

### Protein design

The “PCPcon” program [30, 31] was used to convert alignments of the virus sequences to PCP-consensus sequences. The PCPcon program is related to PCPMer, a program developed to identify PCP-motifs in protein families [47-49]. The PCPs are defined by five quantitative descriptors of each amino acid obtained from multidimensional scaling of 270 physical chemical properties of the amino acids [50]. The program selects for each column in an amino acid alignment that is most similar in their PCPs to all the other naturally occurring in that column [30-32, 51, 52]. This approach to a consensus vaccine differs from that of others (such as [53-55]), which select residues based on frequency of occurrence in a selected group of sequences. Our method automatically reduces the weight of viral sequences with high superficial redundancy (i.e. viral sequences that differ from one another only at a few positions) in the calculation of the conservation of PCPs. We have shown, with examples from four different virus groups, that PCP_con_ peptides and proteins fold in a fashion similar to the wild-type proteins upon which they are based [45, 51], retain function [38, 52], are immunogenic and can stimulate broad spectrum antibodies. For example, a PCPcon antigen (7p8) stimulates antibodies that neutralize representatives of all four dengue serotypes [51].

DGraph is a program that automatically calculates and graphically portrays the pairwise property distances (PD) between unaligned sequences supplied in Fasta format as a text file [56]. Lower PD values indicate a higher sequence similarity. The program is available from Github (https://github.com/bjmnbraun/DGraph).

### Designing peptides from exposed regions of the A domain of E2

We used the previously calculated PCP_con_ sequences of the whole E2 protein of either CHIKV_con_ or AllAV_con_. The latter protein was the PCPcon calculated from sequences of many diverse alphaviruses as described previously [45]. Models of both PCPcon sequences were prepared based on the crystal structure of the CHIKV E2 protein [57]. Based on the anticipated surface-exposed areas of the A domain of these models (from the GETAREA program [58]), we designed and synthesized (within the LANL peptide facility) four peptides from CHIKV_con_ (numbered 1-4) and corresponding ones from the AllAV_con_ (5-8). Details on the sequence, preparation and purification of the peptides are described in supplementary material.

### Peptide Synthesis

All reagents and solvents deployed were of peptide synthesis or biotech grade. Wang solid phase resins were acquired from AdvancedChemTech. All other Fmoc amino acids were purchased from P3Bio with the exception of Fmoc His (tBoc), required for high temperature coupling reactions, which was obtained from CEM. Dimethylformamide (DMF) and the deployed deprotection reagent 20% Pyrrole (prepared as solution in DMF) was obtained from Alfa Aesar. Pyrrole is the suggested Fmoc deprotection reagent by CEM, as elevated reaction temperatures of 105°C in deprotections using piperidine lead to loss of side chain protecting groups. The peptide coupling reagents, Diisopropylcarbodiimide (DIC) and Oxyma Pure (Eythylcyanohyroxyiminoacetate) were acquired in peptide synthesis grade from AKScientific. General reagents such as N,N-diisopropyl ethyl amine (DIPEA), triisopropyl silane (isoPr_3_SiH; TIPS), thioanisole, octaethylenglycol-dithiol and trifluoroacetic acid (TFA) were purchased from Sigma Aldrich, methylene chloride (DCM) and diethylether were obtained from Fisher Scientific. For HPLC purifications, Acetonitrile was acquired from Alfa Aesar and water was purified in-house (deionized, filtered through a Nanopure to 18.2 MΩ*cm resistivity, and UV-sterilized). Mass spectrometry used highest quality (Optima MS grade) solvents purchased from Fisher Scientific.

#### Automated Peptide Synthesis using the CEM Liberty Prime Microwave Peptide Synthesizer

A CEM Liberty Prime microwave peptide synthesizer was used for solid phase synthesis at high temperature (105^0^C). All syntheses were performed at the 0.1 mM scale at the recommended standard instrument chemistry modified to increase drain times from 5 to 10 sec to accommodate for any resin volume increase over the synthesis cycles. For these short peptides, single coupling instrument cycles were used, with achieved average coupling yields for cycle of > 98.5%. The synthesis deploys 65 sec coupling at 105°C, direct addition of the pyrrole to the hot resin and deprotection at 105°C for 45 sec, 20 sec for the three wash steps for a total single couple cycle time of 3 min. To prevent hydrolysis of acid labile side chain protecting groups during the syntheses, 0.1 M DIPEA was added to the Oxyma solution.

#### Deprotection and Removal of the Peptides and Proteins from the Resin

Deprotection used 25 mL of modified “reagent K” mixture: TIPS (1.25 ml/25 ml), thioanisole (0.625 ml/25 ml), octaethyleneglycoldithiol (1.25 mL/25mL) -a less odorous substitute for EDT (ethylendithiol) - and water (1.25 ml/25 ml) in TFA (trifluoroacetic acid). The resin was pretreated with the quencher solution for 5 min, then TFA was added (to a final volume of 25 ml). The deprotections were caried out in 50 mL conical tubes of high-density polypropylene, under a blanket of Argon to prevent side reactions from air for 1.5 hours at room temperature. The solutions were filtered and the filtrate was concentrated to 10 mL. The peptides were then precipitated into ice cold ether and collected by centrifugation.

#### Purification and Analysis of synthetic peptides

Purifications (to> 98% +) were performed on a Waters HPLC preparative workstation with 2545 pump (at 20 ml/min), using a C_18_ reverse phase column (Waters BEH 130, 5 μm, 19×150) and a gradient from water to acetonitrile with 0.1% TFA added. Peaks were collected based on monitoring at 215 nm using a PDA 2998 detector. Combined product fractions were lyophilized, yielding white fluffy solids. Peptides were then analyzed for purity by analytical HPLC on a C_18_ reverse phase column (Waters BEH 130, 5 μm, 4.6×150) with a linear gradient from 96% to 20 % water-acetonitrile with 0.1% TFA in 12 minutes and by mass spectrometry on Thermo LTQ or Thermo Exactive mass spectrometers in ESI+ mode. Additional data on the HPLC conditions and mass spectrometry identification are provided in the compound listing.

### Gene cloning

The gene sequences for the consensus proteins of the B region of the E2 protein, e.g., amino acids 170-268 of CHIKV, were codon-optimized for *E*.*coli* expression, synthesized and cloned into the pCold II plasmid vector in the multicloning site between the Nde1 and Xba1 sites by Biobasic (Toronto, Canada). Each approximately 97 amino acid stretch was preceded by a 14 amino acid segment containing a hexahistidine tag followed by a four amino acid recognition site to allow cleavage to the native protein by Factor Xa.

### Protein expression

Plasmids were transformed into *E. coli* Origami 2 competent cells (Novagen). Transformed cells were stored as glycerol stocks at -80 °C until use. To express protein, a 1.2 mL aliquot was divided among 4x 4 L flasks each containing 1 L of 2x LB broth (20 g/L tryptone (Fisher Bioreagents), 10 g/L yeast extract (Sigma) and 10 g/L NaCl ((Fisher) set to pH 8 with 10 M NaOH. After autoclaving, 1 mM MgSO_4_ (Fisher) 50 µg/mL carbenicillin (Sigma) and 12.5 µg/mL tetracycline (Sigma) were added. Cultures were grown overnight at 37 °C with vigorous agitation (230 rpm). Then 1 L of LB medium containing 50 µg/mL ampicillin was added. Following 1 h of further agitation at 37 °C the flasks were cooled on ice for approximately 15 min, 1 mM IPTG (Goldbio) was added and the cultures were set at 15 °C for 24 h (230 rpm). Cell pellets were collected by 20 min centrifugation at 5000 x g (4 °C).

### Protein isolation

Cells were resuspended in lysis buffer (20 mM Tris/HCl pH 8 (Tris Base, Tris Hydrochloride, Fisher), 750 mM NaCl, 10% glycerol (Fisher) containing 1 mM phenylmethylsulfonyl fluoride, PMSF (MP Biomedical) and protease inhibitor tablets (1 tablet/100 mL, cOmplete Mini EDTA-Free, Roche Diagnostics), 5ml / g wet weight of pellet. Cultures were lysed by freezing and thawing (−80 C freeze followed by thawing in RT water) and sonicating for 10 min (cycle: 30 s on. 30 s off; Virtis, Virsonic 550). Lysate supernatants were clarified by 30 min centrifugation at 15,000 rpm (4 °C).

About 130 ml of clarified cell suspension was combined with 3 mL of TALON^®^ resin (Takara Bio 635503) previously equilibrated in lysis buffer and rocked at 4°C for 1 h. Typically, the treated resin was washed (3x) in 30 mL FB (final buffer: 20 mM Tris/HCl pH 8, 300 mM NaCl, 10% glycerol), poured into a chromatography column (Bio-Rad Poly-Prep^®^), washed additionally with 5 mM imidazole in FB (15 mL) and the column eluted with 5 mL aliquots of FB with 10-150 mM imidazole

The N-terminal His tag was removed by dialyzing (6000-8000 MW cutoff tubing) the TALON^®^ protein containing eluates against FB (20x excess) overnight to remove small contaminants and imidazole. Then CaCl_2_ was added to 2mM and 1 µL (about 1 µg) of Factor Xa (NEB) was added per 50 µg of protein and the samples were left at RT for 20-24h. The samples were passed through a 30 kDa cutoff Amicon^®^ Ultra centrifugal filter unit to remove the Factor Xa and residual high molecular weight contaminants. The flow through was concentrated in a 3 kDa cutoff Amicon^®^ Ultra filter unit (typically from an 20-30 mL vol to 4 mL) and buffer exchanged 3x by adding 10 ml of FB and reconcentrating to remove most of the His tag fragment, residual imidazole and CaCl_2_. The sample was reapplied to a gravity flow 1 mL TALON^®^ column or a TALON^®^ spin column (635601) to remove any remaining N-terminal fragment and uncleaved protein.

### Protein characterization

<u>Protein purity</u> was assessed by polyacrylamide gel electrophoresis (PAGE,) and MALDI-mass spectroscopy. Tricine-SDS-PAGE experiments were performed using a single layer 0.75 mm thick 20% acrylamide/bis-acrylamide gel (using 29:1 Acrylamide/Bisacrylamide, Bio-Rad) run at 120 V constant voltage (Pharmacia LKB, ECPS 3000/150) using a Hoefer gel system and Precision Plus Protein Kaleidoscope standards (Bio-Rad). Protein and peptide concentrations were determined by direct measurement on a Nanodrop 1000 UV-Vis spectrophotometer (Thermo Scientific), using peaks around 280 and 230 nM, respectively. The 260/280 ratio of the purified proteins used for inocula was between 0.6 and 0.7. Protein concentration was also measured by a Coomassie Blue (Bradford) assay using a BSA standard with a SpectraMax M2 plate-reader (Molecular Devices).

<u>Protein molecular masses</u> were measured with a Sciex 5800 MALDI-TOF/TOF mass spectrometer using horse heart cytochrome c external standard (mw 12,384 Da, Sigma) in positive ion mode (m/z range 4000-35000 Da, focal mass 15,000 Da, 2000 laser shots/spectrum). For MALDI analysis, samples in final buffer and external standards were mixed 1:20 (v/v) with 10 mg/mL sinapic acid in 50% water/50% acetonitrile/0.1% formic acid; of which a 1.2 µL sample was dried on a MALDI plate at 37 °C.

<u>Circular dichroism (CD) spectra</u> were collected on a Jasco J-815 spectrometer (DIY=8sec, scan speed 20nM/min, 10 iterations 178-260 nM) and interpreted with the CDSSTR program [59, 60], accessed via the Dichroweb server [61].

<u>Protein dotspot assay</u> was used to characterize the antibody specificity of sera of inoculated rabbits. For dotspots, proteins (1 µl aliquots, 0.25 µg) were dotted onto nitrocellulose. The spots were allowed to dry in air and the membrane strips were blocked with 5% milk in PBS buffer with 0.1% Tween 20 for 5 min. The blot was washed with PBS and developed for 1h at RT with serum samples diluted 1:200 with PBS. The blots were washed with PBS and further developed with goat anti-rabbit IgG-horseradish peroxidase labeled (HRP) (Southern Biotech #4050-05) diluted 1:1000 for 1 h at RT. The membranes were washed and developed with 4-chloronapthol-H_2_O_2_ reagent.

### Protein immunogenicity

Antisera against the consensus antigens EEEV_con_, VEEV_con_, CHIKV_con_ and EVC_con_ were generated in male New Zealand white rabbits (Immunotech, Ramona, CA). Initial inoculation of 200 µg of each purified protein in Freund’s complete adjuvant (final volume, 0.5 ml) was followed after 3 weeks by 2 boosts, 3 weeks apart of 100 µg in Freund’s incomplete adjuvant. Rabbits were bled three weeks after the second and third immunizations.

A variation on this protocol was the addition of 10 µg each of either peptides 1-4 or 5-8 from the A domain of CHIKV_con_ or AllAV, as described above and in supplementary material. Peptides were quantitated based on their absorbance peak around 230 nM and kept frozen at -80° until incorporated into the inoculum with the B region recombinant proteins (which were stored at 4°C until use).

To purify the antibody-containing fraction, 25 ml of serum was combined slowly with 14 ml saturated Ammonium sulfate and the samples left overnight at 4°C to precipitate. They were centrifuged for 30 min at 3500 x g, the supernatant was discarded and the pellet taken up in 5 ml of sterile bi-distilled water. The solubilized pellets were dialyzed (12-14kD cutoff dialysis sacks, Spectrapore) against 3 buffer changes of PBS. The final volume of sample was about 8 ml for an overall concentration factor of about 3 compared to the starting sera.

#### Plaque Reduction Neutralization Test (PRNT)

Sera were tested for neutralizing antibodies as described previously [38, 62]. Sera were inactivated for 30 min at 56°C, then serial dilutions were incubated in a 12-well plate for another hour at 37°C with ∼800 pfu/mL of virus strains: VEEV-TC83 (29 plaques in the negative control), CHIKV181-25 (24 plaques) [63] and EEEV105 (30 plaques). Purified antibody preparations were not heated before diluting for PRNT. Vero cells were infected with sera/virus mixtures, and plaques were counted to determine PRNT_80_ (80% reduction) or PRNT_50_ titers where applicable. For example, if the 1:80 sera dilution was the highest dilution with 80% or fewer plaques than the average in the mock-treated wells, the serum was considered to have a PRNT_80_ of 80. For PRNT against the EEEV related Madariaga virus, MADV76V25343 and the VEEV related Mucambo [64], MUCV BeAn-8, untreated controls had 55 plaques and 50 plaques, respectively.

## Results

### PCPcon proteins represent individual species or several AV

As outlined in Materials and Methods and a previous paper [45], we prepared recombinant B-region protein fragments of the E2 envelope proteins. This surface-exposed region of the “trimeric spike”, near the putative receptor binding site [65], has been shown to mediate binding of neutralizing monoclonal antibodies against VEEV or CHIKV [41-43, 65-67]. We first calculated PCPcon sequences for the E2 protein based on unique amino acid sequences of natural isolates of three diverse alphaviruses: 33 strains of EEEV, 93 strains of VEEV and 194 strains of CHIIKV. As Figure 1A shows, the three PCP_con_ proteins have low sequence identities below 45%. In addition, we produced EVC_con_, based on the PCPcon sequences EEEV_con_, VEEV_con_, and CHIKV_con_, which has around 50% identity to each of the individual sequences it was based on (Fig. 1A) and represents a PCP-mean of the diverse viruses in the PD-graph (Fig. 1B). The PD-graph separates the individual viruses of the EEEV, VEEV and CHIKV complexes in distinct regions. The purified proteins used for inocula were pure according to MS and PAGE (Fig. 2) and have circular dichroism spectra consistent with a properly folded picture (results previously shown and Fig. S2).

**Figure 2.**
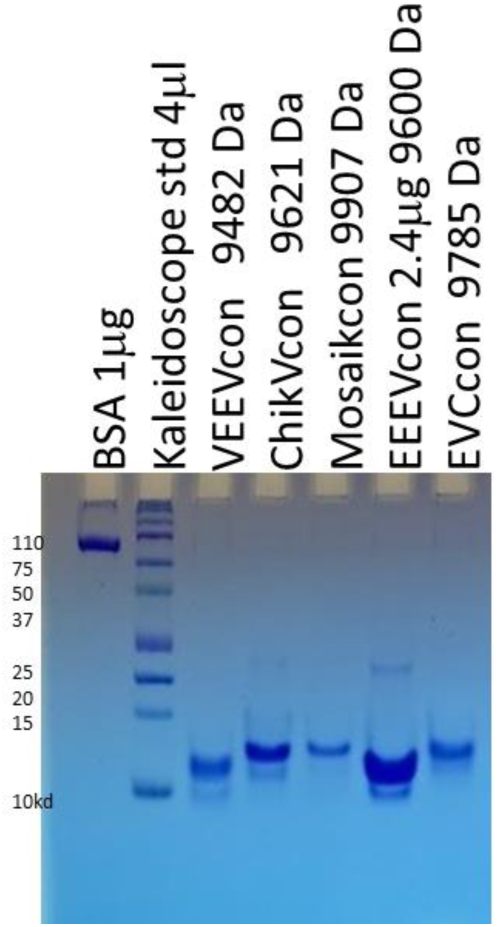
Purified PCPcon protein antigens used for inocula. PAGE (16% gel) of proteins (1 µg per lane, except as indicated) were purified as described and used for inoculations alone or with A region peptides as described in Materials and Methods.

### Inoculation with wild-type PCPcon proteins induces specific antibodies

We previously showed that PCP-consensus sequences, VEEV_con_ and CHIKV_con_, based on unique sequences of the “B” domain of the E2 protein, were recognized specifically by antibodies in sera after infection with either the VEEV-TC83 or CHIKV-Ross strains [45]. Consistent with our previous results, the species-specific proteins, VEEV_con_, EEEV_con_ and CHIKV_con_, induced serum antibodies that recognized only the immunizing protein, and to some extent the EVC_con_ protein (Fig. 3). In contrast, EVC_con_ induced antibodies that recognized all three species-specific proteins. A similar specificity pattern was seen in plaque neutralization tests (PRNT) against prototype viruses of the three AV species (Figure S3, 5). While both EEEV_con_ and VEEV_con_ antisera specifically neutralized their cognate viruses, immunization with the CHIKV_con_ protein alone did not yield CHIKV neutralizing antisera.

**Figure 3:**
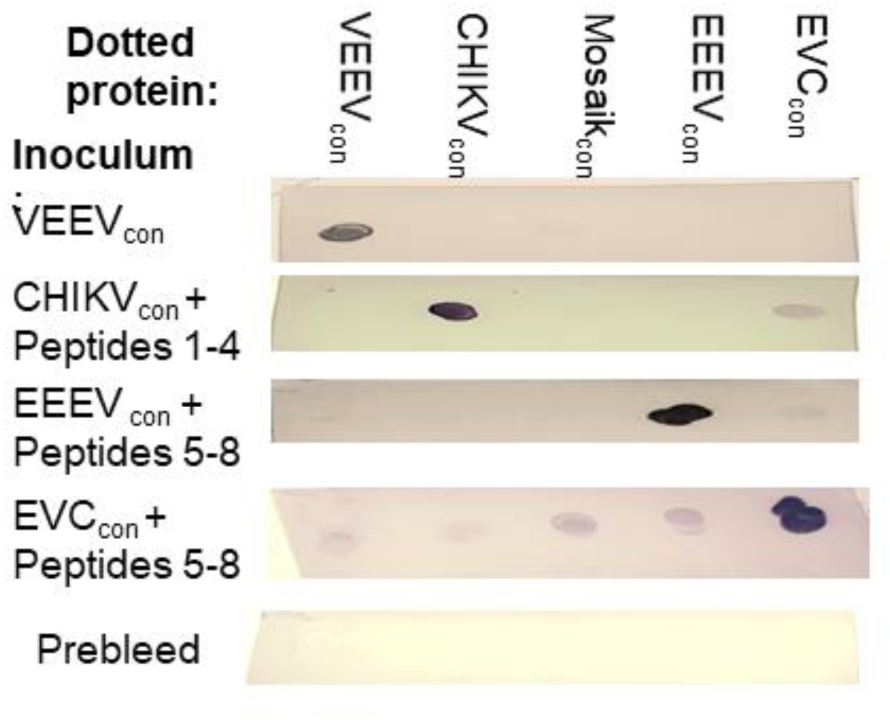
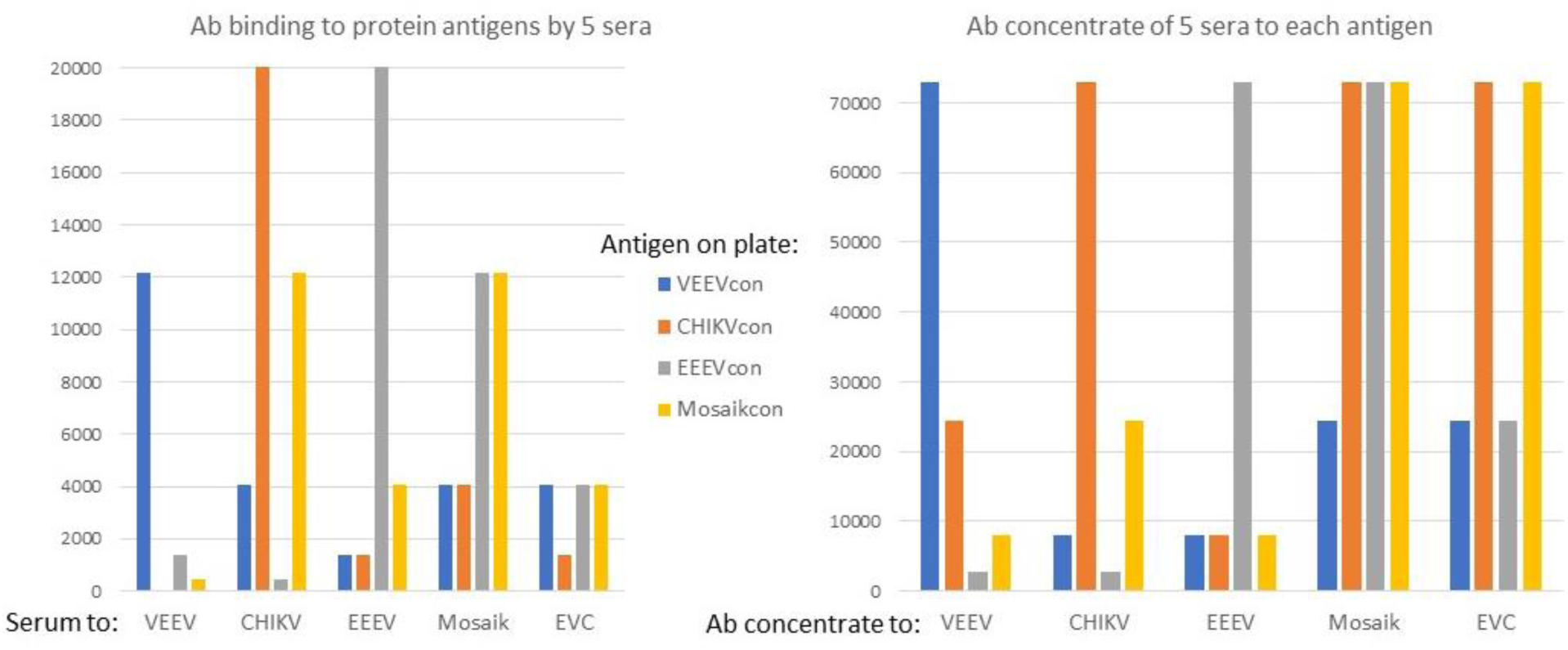
Specificity of serum antibody response to species-specific or consensus proteins after three inoculations. Sera generated in response to VEEV_con_, CHIKV_con_, and EEEV_con_ antigens, calculated based on the ensemble of strains shown in Fig. 1, stimulate specific immune responses to themselves. Serum antibodies generated in response to inoculation with Mosaic2_con_ or EVC_con_ are broad spectrum and recognize all the other proteins. A) Dotspots (0.25 µg each recombinant protein) were developed with the indicated sera. The EVC_con_ protein is also recognized to some extent by antibodies in sera after inoculation with CHIKV_con_ and EEEV_con_, further emphasizing that it presents epitopes common to multiple alphavirus proteins. B) ELISA assays against four recombinant proteins using sera (left, same as used for dotspot) or their antibody concentrates (right) show. Purified recombinant proteins were applied to the 96 well plates and tested against 5 sera or purified antibody concentrates isolated from them. Sera from rabbits inoculated with Mosaik2_con_ or EVC_con_ + peptide 5-8 and their concentrated antibody solutions recognize all four recombinant proteins, while sera generated against the serotype specific antigens VEEVcon, CHIKVcon, EEEVcon, recognize their cognate protein and Mosaik2_con_. Data are given as the half maximum serum dilution to see antibody binding: maximum dilution for the sera (left) is 36,000 fold; for the concentrates (right), it is 72,900. Peptides are described in Fig. 4

### Peptides from surface exposed areas of the A domain were needed to obtain CHIKV neutralization

As previous reports had indicated that additional areas of the E2 protein were needed to obtain CHIKV neutralization [68], we designed a series of peptides from the surface-exposed areas of the CHIKV_con_ and the aligned sequence of the AllAV_con_ proteins, using 3D models of their predicted structure (Fig. 4). The sequences of these peptides and their purification details are defined in the Table in Fig. 4. We obtained relatively weak neutralizing activity against CHIKV by adding peptides 1-4 from surface-exposed areas of the A domain to the recombinant CHIKV_con_ B domain in the inocula. Adding PCPcon peptides 5-8 from the corresponding areas of the AllAV-A domain to the species-specific proteins did not enhance the ability of sera to recognize the spike domain from the other viruses (Figure 3). However, combining the EEEV_con_ protein with peptides 5-8 from an AllAV calculated sequence slightly broadened its ability to generate sera that neutralized CHIKV and VEEV, which was particularly clear from the purified antibody fraction (last lanes of Fig. 5).

**Figure 4.**
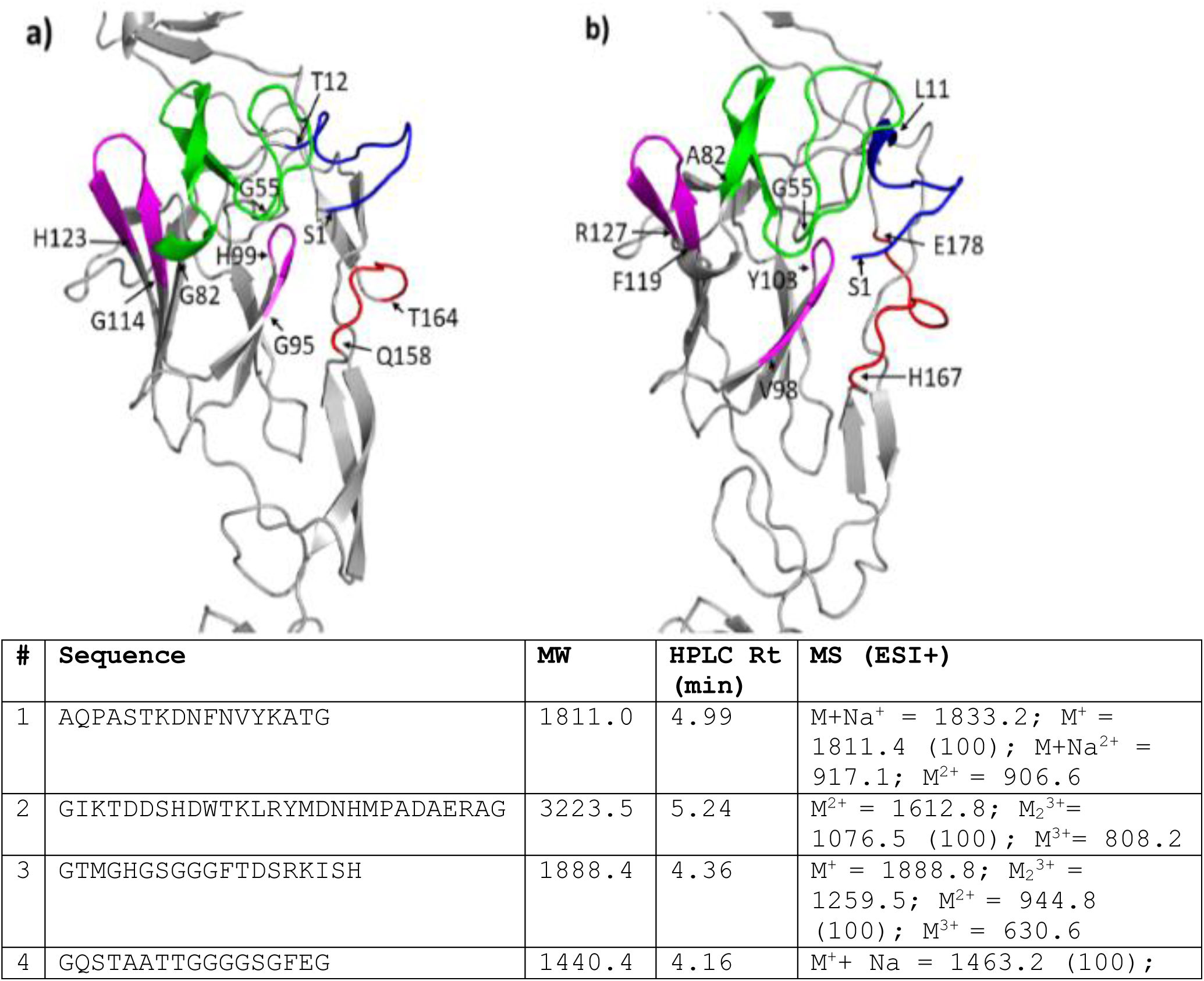
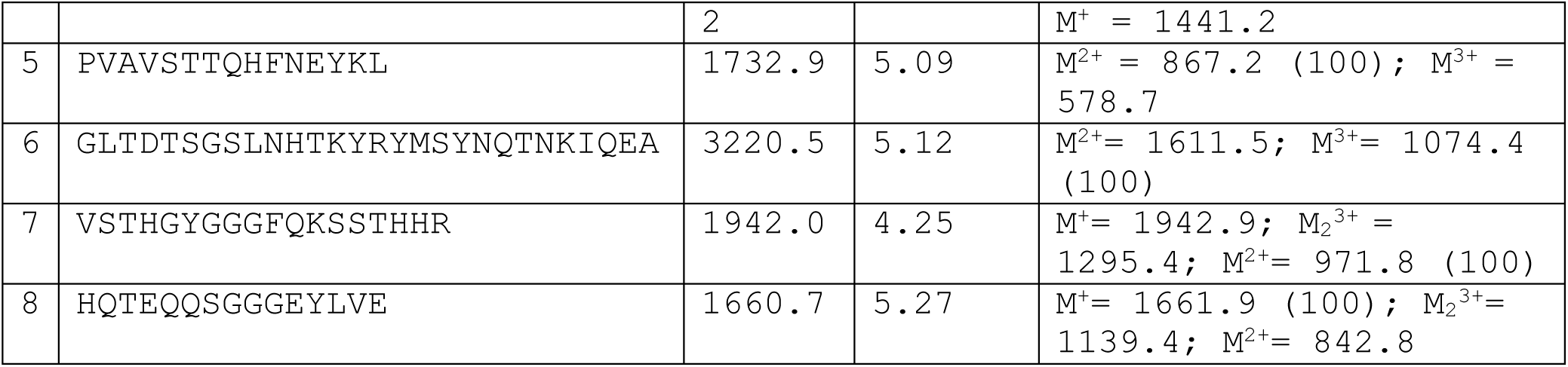
Peptides 1-4 were derived from a PCPcon of the E2-A domain of CHIV strains, Peptides 5-8 were derived from were selected based on alignment of the E2-A domain of CHIKVcon and AllAVcon sequence (see supplementary material), based on 24 different AV species. The AllAV B-domain was the basis for Mosaikcon, which was modified to contain known neutralizing epitopes of VEEV and CHIKV[45]. A,B: Surface-exposed peptides mapped on models of the E2-A-domain of CHIKV_con_ (a) and AllAV_con_. Table: sequence and purification details of the peptides.

**Figure 5:**
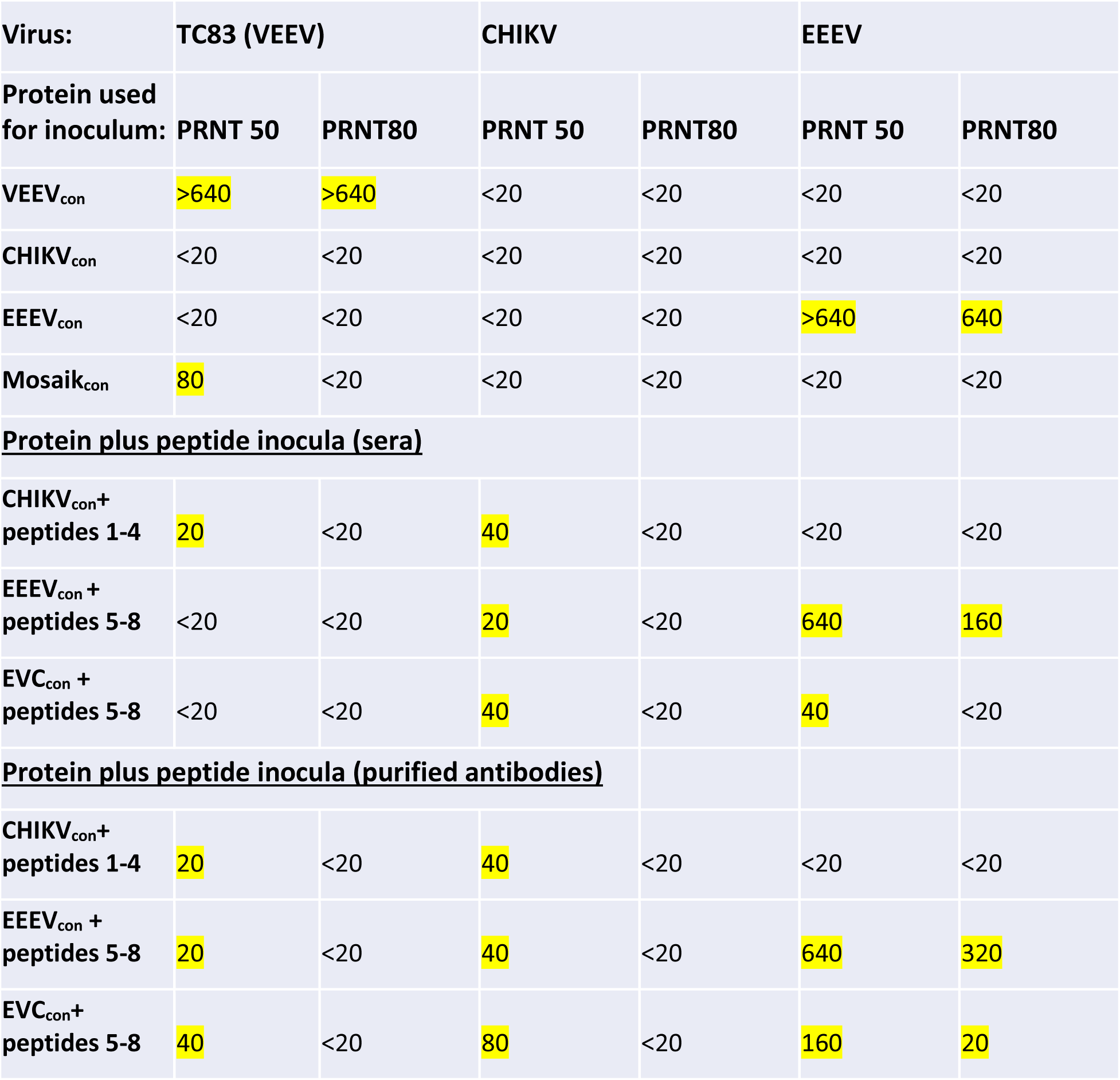
Plaque neutralization (PRNT_50_ and PRNT_80_) titers of antibodies in sera from rabbits inoculated with recombinant PCP_con_ antigens. Sera were diluted from 20-to 640-fold and numbers indicate maximum fold serum dilution to retain 50% or 80% (PRNT_50_/PRNT_80_) reduction in virus plaques. Results for purified antibodies (which gives about a 3x concentration factor) are also shown for EEEV_con_ and EVC_con_ (which were supplemented with A region peptides in the 3^rd^ inoculum as shown in Figure 4).

The antisera showed similar selectivity in neutralizing different viruses from within the VEE and EEE antigenic complexes (Fig. 1B). As anticipated from assays done with antisera generated in cotton rats to EEEV [1], our EEEV_con_ antisera neutralized MADV with a PRNT_50_ of 80 and a PRNT_80_ of 40, or about 4-fold lower than that seen for EEEV (Fig. S3). Similarly, the VEEV_con_ antisera only neutralized Mucambo virus (MUCV; another species in the VEE antigenic complex) with a PRNT_50_ of 20 (Fig. S3). When our VEEV_con_ protein sequence was incorporated into a whole live-attenuated vaccine, PRNT activity in the sera of inoculated mice were relatively low. However, the vaccine prevented lethality with MUCV and (partially) MADV [38].

### Purely computational, EVCcon generates a broad spectrum antibody response

In contrast to the species specificity of EEEV_con_, VEEV_con_, and CHIKV_con_, antisera generated to the EVC_con_ protein whose sequence was calculated based on all three of these, plus peptides 5-8, recognized all the other protein antigens, regardless of species or absolute identity in both dotspots and ELISA (Fig. 3). As Fig. 5 shows, the combined inoculum generated a serum antibody ensemble that neutralized viruses from each of the 3 antigenic complexes.

## Discussion

These results show that it is possible to computationally design a protein/peptide inoculum to generate neutralizing antibodies against 3 different alphavirus species with over 60% amino acid sequence diversity. We also show that PCP_con_ alphavirus proteins can be designed to generate either a species-specific or a promiscuous, protective immune response. Comparison of the alphavirus family revealed that the sequences of EEEV were intermediate in amino acid ID to both the VEEV-and the CHIKV-related AV groups (Fig. 1). We also produced EVC_con_, a protein that represents the average PCPs of VEEV_con_, CHIKV_con_ and EEEV_con_ and is intermediate in PCPs to all alphavirus sequences (Fig. 1B). Immunizations with VEEV_con_, EEEV_con_ or CHIKV_con_ generated species-specific neutralizing antibodies with limited cross reactivity (Fig. 3, 5). The most cross-reactive sera were obtained by inoculation with EVC_con_, a completely computational protein, coupled with peptides from an AllAV E2 sequence calculated from 24 different AV (Fig. 4).

Our results imply that specificity cannot lie in pure sequence identity, as both EEEV_con_ and EVC_con_ have similar overall amino acid identity to VEEV_con_ and CHIKV_con_. EEEV_con_ retained species specificity and generated antibodies that primarily recognized and neutralized EEEV or its South American relative in the EEE complex, Madariaga virus (MADV). In contrast, EVC_con_ stimulated a promiscuous, *polyclonal*, broad-spectrum response that neutralized all three viruses. Adding peptides representing surface-exposed regions of the A domain of the CHIKV_con_ E2 sequence, analogous to the approach of Weber [68], was necessary to obtain CHIKV-neutralizing antisera. We also showed that adding analogous peptides representing the average PCPs of the A regions, calculated for 24 different alphaviruses (AllAV, [45]), enhanced and broadened the neutralizing activity of the sera induced by both EEEV_con_ and EVC_con_. These results illustrate how the PCPcon approach can enhance the design of vaccines to generate immunity against diverse viral species.

There are many confusing examples of the degree of protection provided by AV vaccines. For example, EEEV and its South American relative MADV are very closely related (Fig. S1). Although these viruses cross react serologically, the sera of cotton rats infected with EEEV give only 4-fold reduced plaque reduction neutralization titers (PRNT) against South and Central American MADV isolates compared with titers against N. American EEEV [1]. Yet EEEV vaccination can give limited protection against the much more divergent VEEV and even CHIKV. We have obtained similar results with sera of rabbits inoculated with the EEEV_con_ E2 protein, which gave robust PRNT_50_ values of >640 for EEEV, but only 80 against MADV (Fig. S3). Further, VEEV/IRES live-attenuated vaccines provided some protection against MADV [38]. In our experiments, while the EEEV_con_ protein has only slightly less identity to VEEV_con_ or CHIKV_con_ than does EVC_con_ (Fig. 1A), it generates a species-specific response that is only slightly broadened by adding peptides from the AllAV-A domain of the E2 protein. While EEEV’s PD values place it in a discrete group with MADV, the EVC_con_ protein lies in the middle of the PD graph (Fig. 1B), showing it represents properties central to all alphaviruses.

### Distinguishing species specificity from promiscuity in antigen design

In vivo, as viruses spread from cell to cell and reach high titers in tissues, the surface of virus particles that survive neutralization and antibody-and complement-driven lysis move further away from the initial sequence ensemble of the infecting virus [69]. Cell culture passaging of a single strain in the presence of high affinity monoclonal antibodies gives rise to viral variants that “escape” neutralization. Even single amino acid changes in a peptide or protein can prevent binding to mAbs [70].

These results have important implications for the design of vaccines and therapeutics against rapidly mutating viruses and further emphasize that the polyclonal antibody response to discrete antigens can give broad spectrum protection. The PCPcon approach takes advantage of the promiscuity built into the immune response. Polyclonal antibodies can have overlapping binding sites even to the same peptide epitope [71, 72] and may recognize multiple similar epitopes. For example, in other experiments, we showed that polyclonal antibodies from sera of SARS-CoV-2-infected patients recognize the spike protein of the SARS-2003 virus, despite <80% sequence identity [73]. In the continuation of this alphavirus work, we will analyze the epitope specificity of the antibody ensemble produced in response to our antigens, with the goal of elevating the protective effect of the promiscuous immunogens to that of the species-specific ones against each AV.

## In conclusion

these experiments show that protein/peptide, broad spectrum antigens designed to represent the common PCPs of the most exposed areas of the E2 protein can provide some degree of protection against AV with less than 60% amino acid sequence variability. Such antigens can greatly simplify the design of vaccines and eventually therapeutic antibodies against evolving pathogenic viruses.

## Acknowledgements

This work was supported by NIH grants R01 AI137332 and R24 AI120942. Peptide syntheses at LANL were supported by LDRD ER funding. We thank Prof. Thomas Ksiazek for advice on antibody purification and other members of the World Reference Center for Emerging Viruses and Arboviruses and the Animal Resource Center at UTMB for their help.

