## SupplementaryFig1-4 for "Peptide and protein alphavirus antigens for broad spectrum vaccine design"

Figure S1: **PD-graph of the E2 protein of EEEV strains from the US** (which cluster (below right), regardless of host or year, around our $EEEV_{con}$) and related isolates from South & Central America. Cutoff PD <4 means all strains are very closely related (compare to the PD-graph in the text, Fig. 1, where the cutoff is 14).

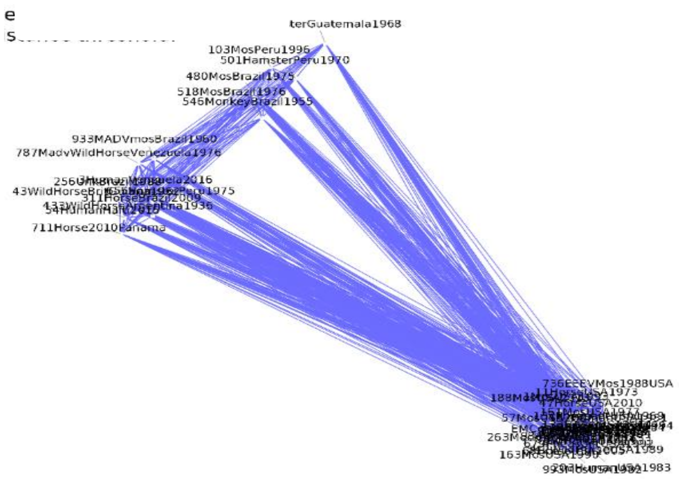


Figure S2: PAGE (Coomassie blue strain) and Maldi Mass spectroscopy data (anticipated MM, 9785, consistent with one or two (4904 peak) sodium atoms) for the $EVC_{con}$ used for rabbit inoculation.

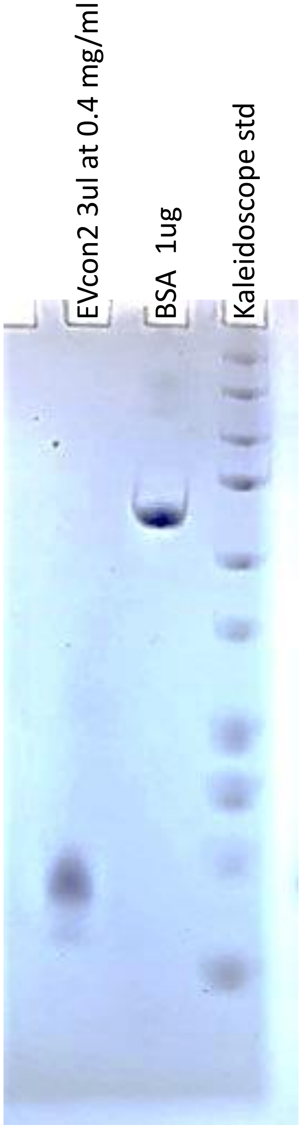


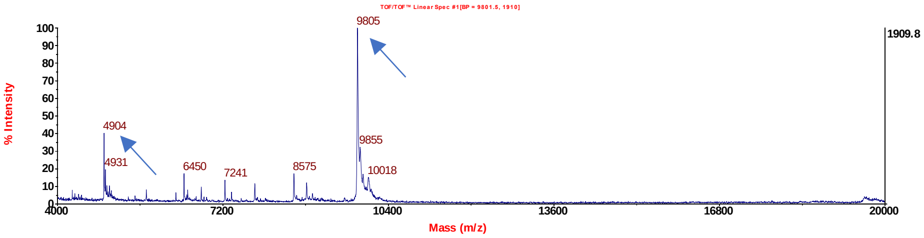

Figure S3: PRNT data for the sera of rabbits inoculated with PCPcon proteins (see Baker et al. 2020) and tested against 5 different viruses. Note no serum was protective against CHIKV unless A region peptides were added to the inoculum (see Fig. 4 in the main text). The sera were collected 6 months after rabbits were inoculated 3x with the indicated protein (initial in FCA, two booster in FIA) and then 3 weeks after a 4th booster in FIA.

| | | TC83 | | MUCV | | EEEV | | MADV | |
|---|---|---|---|---|---|---|---|---|---|
| **Sample #** | **Sample** | PRNT 80 | PRNT 50 | PRNT 80 | PRNT 50 | PRNT 80 | PRNT 50 | PRNT 80 | PRNT 50 |
| 1 | VEEcon6mos | >640 | >640 | <20 | 1:20 | <20 | <20 | <20 | <20 |
| 2 | VEEcon4x | >640 | >640 | <20 | <20 | <20 | <20 | <20 | <20 |
| 3 | CHIKcon6mos | <20 | <20 | <20 | <20 | <20 | <20 | <20 | <20 |
| 4 | CHIKcon4x | <20 | <20 | <20 | <20 | <20 | <20 | <20 | <20 |
| 5 | MOSAIKcon 6mos | <20 | <20 | <20 | <20 | <20 | <20 | <20 | <20 |
| 6 | MOSAIKcon4x | <20 | 80 | <20 | <20 | <20 | <20 | <20 | <20 |
| 7 | EEEVcon prebleed | <20 | <20 | <20 | <20 | 20 | 40 | <20 | <20 |
| 8 | EEEVcon 2x | <20 | <20 | <20 | <20 | 160 | >640 | 40 | 80 |
| 9 | EEEVcon 3x | <20 | <20 | <20 | <20 | 640 | >640 | 40 | 80 |
| 10 | EEEVcon 4x | <20 | <20 | <20 | <20 | 640 | >640 | 40 | 80 |
| 11 | MIAFControl + (for the respective viruses) | >640 | >640 | 80 | 320 | 160 | >640 | nd | nd |
| 12 | Neg.Control -PBS | <20 | <20 | <20 | <20 | <20 | <20 | <20 | <20 |

Nd: not done

Figure S4: Peptides from the E2-A domain of CHIKV and AllAV were obtained from the following alignment (highlighted areas were surface exposed in the CHIKV structure):

The A-region of the E2 protein used for peptide design:

```
E2CHIKV      ---------SVKDNFNVYKATRPYLAHCPDCGEGHSCHSPVALERIRNEATDGTLKIQVS    51
E2all        AVSTSPVAVSTTQHFNEYKLTRPYVAHCSRCAHGHSCHSPIAIEQVQSDSHDGYVRIQTS    60
E2VEEV       ---------STEELFKEYKLTRPYMARCVRCAVG-SCHSPIAIEAVKSDGHDGYVRLQTS    50
                      *. : *: ** ****:*:*  *. * *****:*:* ::.:. ** :::*.*

E2CHIKV      LQIGIKTDD-SHDWTKLRYMDNHMPADAERAGLFVRT---SAPCTITGTMGHFILARCPK    107
E2all        AQFGLTDTSGSLNHTKYRYMSYNQTNKIQEATLHQVTVHTSQPCHVVSTHGYFLLARCPP    120
E2VEEV       SQYGLDSSG-N---LKGRTMRYNMHGTIEEIPLHQVSLHTSRPCHIVDGHGYFLLARCPA    106
              * *:   . .    * * *  :     :.     : ::** ** :..  *:*:*****

E2CHIKV      GETLTVGFTDSRKISHSCTHPFHHDPPVIGREKFH---SRPQHGK--ELPCSTYVQSTAA    162
E2all        GDTITVSFQKSSTHHRTCTVQYKVKFQPVGREKYTRERHPPQHGIELTKPCQVYTHQTEQ    180
E2VEEV       GDSITMEFKKDSVT-HSCSVPYEVKFNPVGRELYT---HPPEHGA--EHPCQVYAHDAQQ    160
             *:::*: * ..    ::*:  :. .   :*** :      *:**     **..*.:.:

E2CHIKV      T---AEEIEVHMP
E2all        QSEYLVEMHDAMP
E2VEEV       RGAYVE---MHLP
```
